# The nucleolar DExD/H protein Hel66 is involved in ribosome biogenesis in *Trypanosoma brucei*

**DOI:** 10.1101/2021.06.11.448066

**Authors:** Majeed Bakari-Soale, Nonso Josephat Ikenga, Marion Scheibe, Falk Butter, Nicola G. Jones, Susanne Kramer, Markus Engstler

## Abstract

The biosynthesis of ribosomes is a complex cellular process involving ribosomal RNA, ribosomal proteins and several further trans-acting factors. DExD/H box proteins constitute the largest family of trans-acting protein factors involved in this process. Several members of this protein family have been directly implicated in ribosome biogenesis in yeast. In trypanosomes, ribosome biogenesis differs in several features from the process described in yeast. Here, we have identified the DExD/H box helicase Hel66 as being involved in ribosome biogenesis. The protein is unique to Kinetoplastida, localises to the nucleolus and its depletion via RNAi caused a severe growth defect. Loss of the protein resulted in a decrease of global translation and accumulation of rRNA processing intermediates for both the small and large ribosomal subunits. Only a few factors involved in trypanosome rRNA biogenesis have been described so far and our findings contribute to gaining a more comprehensive picture of this essential process.

## INTRODUCTION

Ribosomes are the molecular machines for translation in all organisms. Eukaryotic ribosomes consist of the small 40S subunit, which contains more than 30 proteins as well as the single 18S rRNA and the large 60S subunit which harbours the 5S, 5.8S and 25/28S rRNA and more than 40 proteins. The process of ribosome biogenesis, which starts in the nucleolus and ends in the cytoplasm, is extremely complex and energy-consuming and involves the combined action of small nucleolar RNAs and several protein factors such as endonucleases, exonucleases ATPases, GTPases, and helicases ^1–4^.

RNA helicases mediate structural remodelling of maturing ribosomes ^5^. The enzymes use ATP/NTP to bind and remodel RNA and ribonucleoprotein (RNP) complexes, but some RNA helicases also act as chaperones promoting the rearrangement and formation of optimal RNA secondary structures ^6,7^. In the context of ribosome maturation, RNA helicases of the DExD/H protein family are of particular interest. Most DExD/H proteins belong to the SF2 superfamily of helicases and are highly conserved in nature from viruses and bacteria to humans ^7^. They contain the highly conserved helicase core domain characteristic of the SF2 helicases with the name-giving DExD/H (where x can be any amino acid) in motif II (Walker B motif) ^8^. A large number of RNA helicases have been shown to be multifunctional but most DExD/H helicases are highly specific to their substrate and can rarely be complemented with other helicases. ^7^.

The process of ribosome biogenesis is best characterised in the yeast model system ^9,10^. It begins in the nucleolus with the transcription of a long 35S rRNA primary transcript that undergoes several cleavages and trimmings to produce mature 18S, 5.8S and 25/28S rRNAs. The precursor of the 5S rRNA is independently transcribed by RNA polymerase III in the nucleoplasm. Several trans-acting factors including snoRNAs and RNA helicases are involved in the maturation process. In fact, nineteen out of 37 DExD/H RNA helicases have been implicated in the synthesis of ribosomes in yeast ^11^: the RNA helicases Dbp4, Dbp8, Dhr1, Dhr2, Fal1, Rok1 and Rrp3 are involved in biogenesis of the small subunit (SSU), Drs1, Dbp2, Dbp3, Dbp6, Dbp7, Dbp9, Dbp10, Mtr4, Mak5 and Spb4 participate in the synthesis of large subunit (LSU) precursors while PrP43 and Has1 contribute to both LSU and SSU biogenesis (reviewed in ^5^).

*Trypanosoma brucei* is a flagellated protozoan parasite responsible for African Sleeping sickness and the related cattle disease Nagana. It undergoes several differentiation events during its complex life cycle, shuttling between the mammalian host (bloodstream stages) and the tsetse fly insect vector (several stages, including the procyclic stage that is dominant in the fly midgut); bloodstream and procyclic stages can be cultured *in vitro*. The parasite belongs to the Discoba clade and is highly divergent from the opisthokonts which belong to one of the major eukaryotic kingdoms ^12^. *T. brucei* therefore separated from the major eukaryotic lineage early in evolution and possesses several unique features in its biology including its rRNA metabolism ^13–15^. Most strikingly, the 25/28S rRNA of trypanosomes is processed into six further fragments, two larger ones and four small ones ^16–19^. Trypanosome homologues to proteins involved in ribosome biogenesis in yeast were identified by homology ^14^ and more recently also by purifying early, middle and late processomes of the ribosomal subunits from the related Kinetoplastida *Leishmania tarentolae* ^15^. These include several homologues to DEAD box RNA helicases.

Only few trypanosome proteins have been experimentally shown to participate in rRNA processing or ribosome biogenesis. Three trypanosome-unique nucleolar proteins, presumably belonging to the same complex, are involved in the biogenesis of the large ribosomal subunit, namely the RNA binding proteins p34/p37 and NOP44/46 and the GTP-binding protein NOG1 ^20–24^. The nucleolar Pumilio-domain proteins PUF7 is directly or indirectly required for the initial cleavage of the large precursor rRNA ^25^. The depletion of PUF7, PUF10, NRG1 (nucleolar regulator of GPEET1) and the trypanosome homologue to the eukaryotic ribosome biogenesis protein 1 (ERB1/BOP1 in yeast/human) causes a reduction in 5.8S rRNA and its precursor ^26^. Depletion of the exoribonuclease XRNE causes accumulation of aberrant 18S and 5.8 S rRNAs ^27^. TbPNO1 and TbNOB1 are involved in the specific cleavage of the 3’ end of the 18S rRNA ^28^. TbUTP10 is needed for pre 18S rRNA processing ^29^. In addition, depletion of components of the nuclear RNA surveillance system, the TRAMP complex protein Mtr4 or of proteins from the exosome, cause accumulation of rRNA precursors ^30–32^.

The *T. brucei* genome has about 51 RNA helicases of the DExD/H protein family ^33,34^. Although the DExD/H proteins play important roles in mRNA metabolism, only a few of them have been characterised so far. Here we show that the DExD/H RNA helicase Hel66 is an essential nucleolar protein involved in the processing of rRNAs of both ribosomal subunits.

## RESULTS

### Identification and sequence analysis of Hel66

The protein with the gene ID Tb927.10.1720 was found in a screen for proteins interacting with a conserved motif within the 3’ UTR of the mRNA encoding the variant surface glycoprotein (VSG), the 16mer motif. VSG is the major cell surface protein of bloodstream form trypanosomes and the 16mer motif in its mRNA is essential for mRNA stability ^35,36^ and highly conserved among all VSG isoforms in *T. brucei*. Tb927.10.1720 was the only protein enriched when using the intact 3’ UTR as bait, but not when using three control RNAs (the scrambled 16mer and 8mer, the reverse complement of the scrambled 16mer and 8mer and the reverse complement of the 16mer) (Supplementary Figure S1). However, electrophoretic mobility shift assays failed to demonstrate a direct interaction between a recombinant form of the protein (GST-tagged) and the 16mer RNA (Supplementary Figure S2, S3).

Tb927.10.1720 is annotated as a putative ATP-dependent DEAD/H RNA helicase in the TriTrypDB genome database (https://tritrypdb.org/tritrypdb). Analysis of the protein sequence shows that the motifs characteristic of this class of helicases are highly conserved (Figure 1A, 1B). The walker B motif (DEAD box) has a valine residue at position 3, classifying the enzyme as a DExD helicase ^7^. Tb927.10.1720 encodes an approximately 66 kDa protein conserved across all Kinetoplastida. We will therefore refer to this DExD helicase as Hel66 in this study. There are about 51 other predicted proteins belonging to the DExD/H protein family in *T. brucei* ^33,34,37^. BLAST (Basic Local Alignment Search Tool) analysis indicates that Hel66 is unique to trypanosomatids. The closest BLAST hit outside the Kinetoplastida is DDX21 in *Asarcornis scutulata* (E-value: 2 ×10^−33^, percentage identity: 27.51%, Query cover: 69%), but a reverse BLAST did not return Hel66 as the top hit.

**Figure 1:**
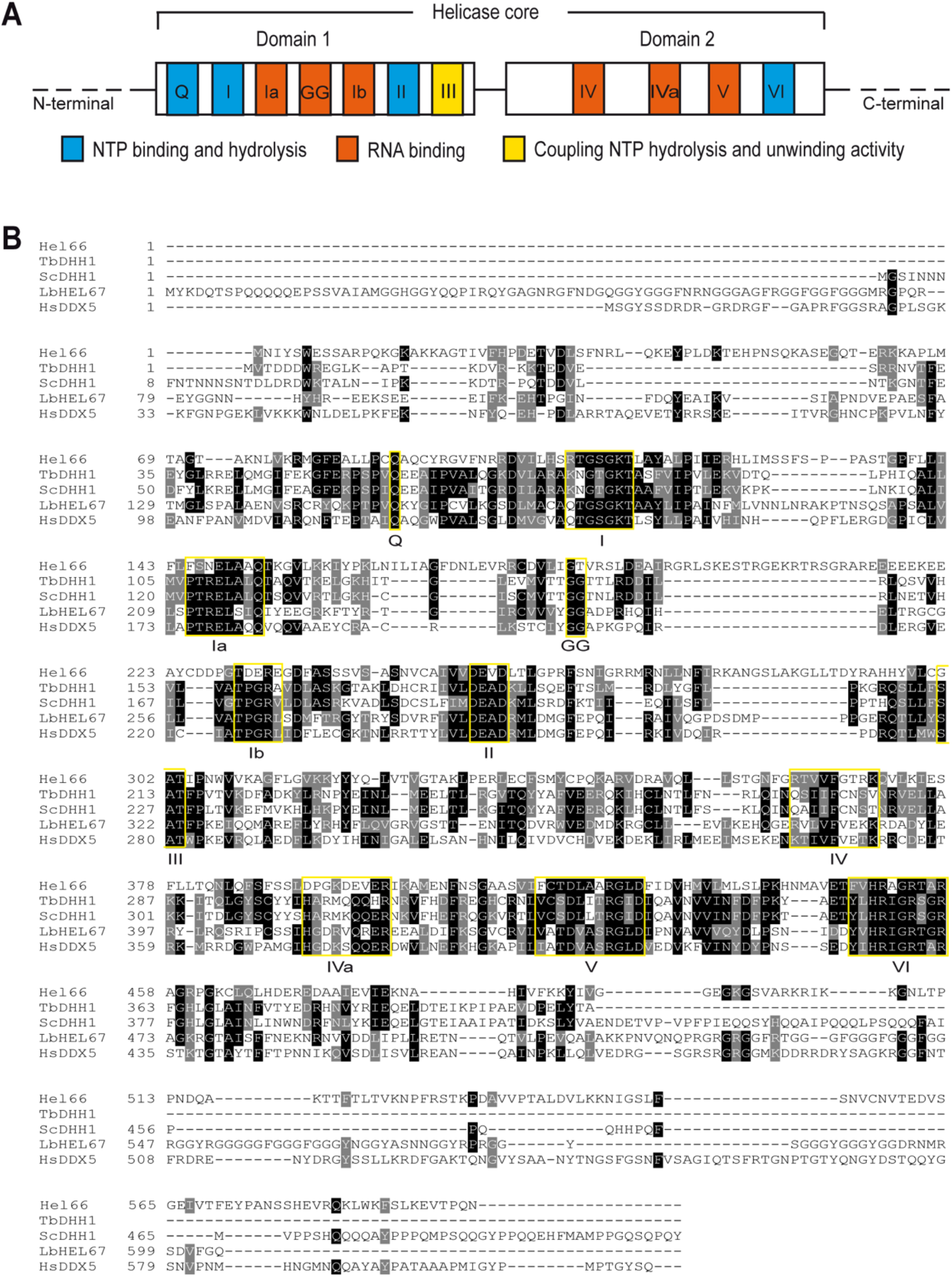
Conserved motifs of DExD/H box proteins present in Hel66. (A) Schematic of the characteristic domains and motifs of DExD/H box proteins. (B) Alignment of Hel66 to previously characterised DEAD box helicase proteins of *Trypanosoma brucei* (TbDHH1), *Saccharomyces cerevisiae* (ScDHH1), *Leishmania braziliensis* (LbHEL67) and *Homo sapiens* (HsDDX5). The characteristic motifs of the DExD/H family are shown in yellow boxes. Black and grey shading indicates identical and similar residues, respectively. The alignment was carried out using Clustal Omega (https://www.ebi.ac.uk/Tools/msa/clustalo/) and formatted for visualisation using BoxShade (https://embnet.vital-it.ch/software/BOX_form.html).

### Hel66 localises to the nucleolus

Hel66 has been previously identified in the nuclear proteome of *T. brucei* ^38^ and the genome wide localisation project TrypTag ^39^ detected the protein in the nucleolus of procyclic-stage trypanosomes. In order to determine the localisation of Hel66 in bloodstream form *T. brucei* cells, we tagged the protein *in situ* at the C-terminus with mNeonGreen. The tagged protein localised to the nucleolus (region of the nucleus with less intense DAPI staining), which is in agreement with the earlier observation in procyclic cells (Figure 2, Supplementary Figure S4).

**Figure 2:**
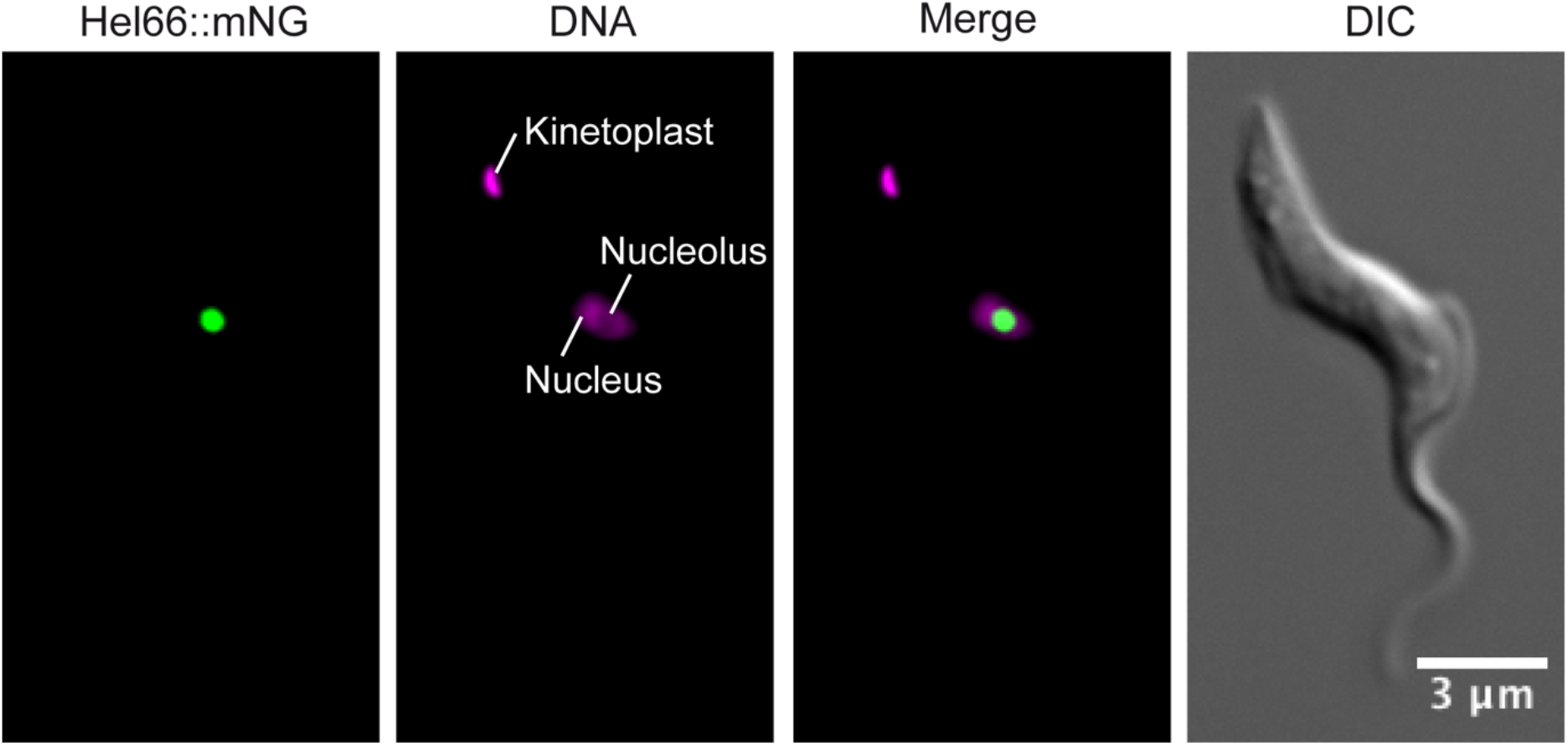
Localisation of Hel66 in BSF *Trypanosoma brucei*. Trypanosomes expressing endogenously tagged Hel66 (Hel66::mNeonGreen) were fixed and imaged. The mNeonGreen signal is concentrated in the nucleolus, the region of the nucleus with less intense DAPI stain. Note that in Kinetoplastida, DAPI also stains the name-giving kinetoplast, the DNA of the single mitochondrion. The images presented in this figure have been deconvolved, the raw images are shown in Supplementary Figure S4.

### Depletion of Hel66 causes stalled growth and perturbation in the cell cycle

The function of Hel66 was further investigated by inducible RNAi, in a cell line expressing endogenously tagged Hel66 (Hel66-HA). Upon RNAi induction, the levels of Hel66-HA decreased to about 20% within the first 24 h and further decreased to less than 10% after 48 h of RNAi induction (Supplementary Figure S5A, S5B). Depletion of Hel66 resulted in stalled growth starting around 24 h after RNAi induction (Figure 3A). In trypanosomes, cells can be assigned to a specific cell cycle stage by counting the number of kinetoplasts (the DNA of the single mitochondrion, K) and the number of nuclei (N) for each cell. As the kinetoplast divides first, a typical (non-synchronised) culture contains mostly 1K1N cells and a smaller number of 2K1N and 2K2N cells. Upon 48 hours of Hel66 depletion, there was a minor increase in the number of cells in the pre-cytokinesis stage (2K2N) and cells with aberrant NK configuration (e.g. multiple nuclei and kinetoplasts) (Figure 3B; Supplementary Figure S5C). These changes in NK configurations suggested a cytokinesis defect but they occurred too late to indicate a direct function of Hel66 in cell cycle regulation.

**Figure 3:**
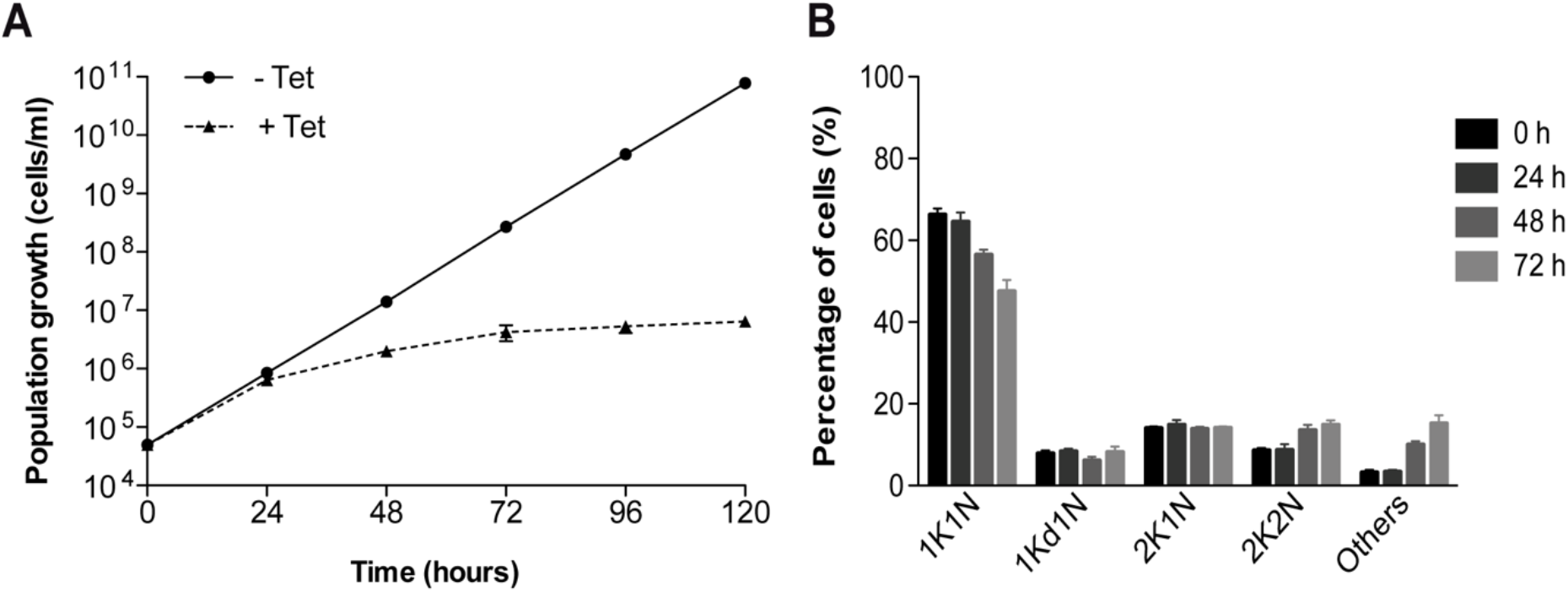
Hel66 is essential in BSF *T. brucei*. (A) Growth curves showing cumulative cell numbers of uninduced (-Tet) and induced (+Tet) Hel66-RNAi cells. Data are averages from three clonal cell lines with error bars representing means ± standard error of the mean (SEM). (B) Cell cycle analysis at different time points during Hel66 depletion. Cells were stained with DAPI and classified to the different cell cycle stages according to the number of nuclei (N) and kinetoplasts (K). As the kinetoplast divides prior to the nucleus, 1K1N, 2K1N and 2K2N stages corresponding to different cell cycle states can be distinguished. 1Kd1N is a precursor stage of 2K1N cells, with the kinetoplast in division. 200 cells were analysed at each timepoint in three clonal cell lines and plotted as mean percentages ± SEM.

As Hel66 was identified in a pulldown assay using a motif of the VSG 3’ UTR RNA sequence as a bait, we investigated whether the protein plays a role in regulating *VSG* mRNA and/or protein levels using RNA dot blots and western blots, respectively. There were no significant changes in *VSG* mRNA or VSG protein levels after 48 h of RNAi mediated Hel66 depletion (Supplementary Figure S6A, S6B). Our data therefore indicates that Hel66 is not a direct interaction partner of the 16mer RNA and has no direct impact on VSG expression. The original interaction between Hel66 and the VSG 3’ UTR was therefore probably unspecific.

### Loss of Hel66 interferes with normal rRNA processing

Many proteins of the DExD/H family are involved in ribosome biogenesis ^30,40–42^. The localisation of Hel66 to the nucleolus, the primary site of ribosome biogenesis, indicates such a function for Hel66 too. We therefore investigated the effect of Hel66 depletion on rRNA processing. Like all eukaryotes, trypanosomes co-transcribe the precursors for the 18S rRNA, the 5.8S rRNA and the 25/28S rRNA as one large precursor transcript (9.2 kb) (Figure 4A). This transcript is initially cleaved into two fragments, the 3.4 kb and the 5.8 kb intermediate. The 3.4 kb intermediate is the precursor for the 18S rRNA. The 5.8 kb intermediate is further processed into the 5.8S rRNA and six further rRNAs that are the trypanosome’s equivalent to the 25/28S rRNA ^16,17^. rRNA processing intermediates were detected by northern blotting using three probes (pre-18S, ITS2 and ITS3) that specifically hybridise to *T. brucei* pre-rRNA, as described by ^13,23,27^ (Figure 4A).

**Figure 4:**
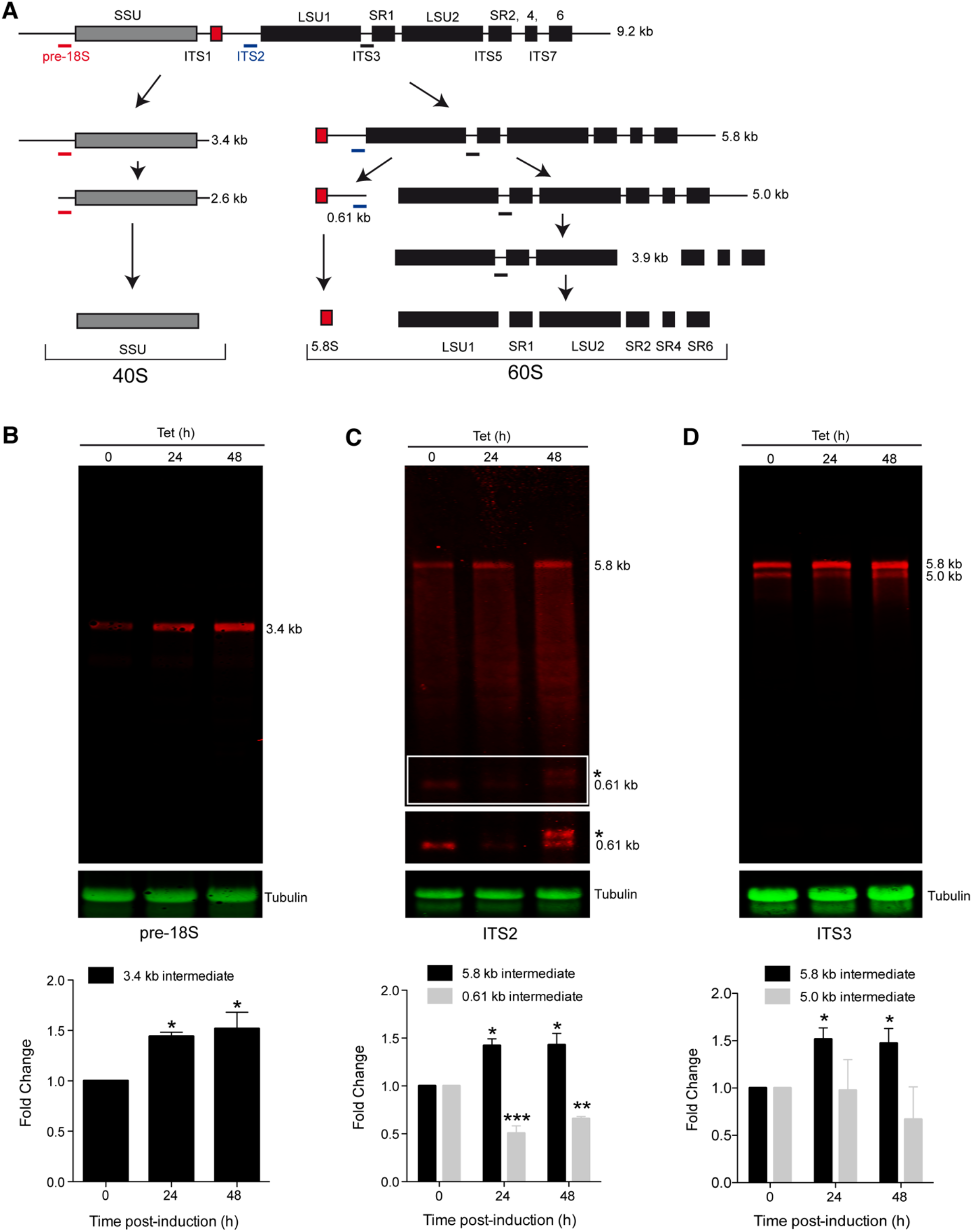
Knockdown of Hel66 results in accumulation of rRNA processing intermediates. (A) Schematic of the rRNA processing pathway in trypanosomes. Probe positions are indicated as thick lines in red, blue and black for the pre-18S, ITS2 and ITS3 probes, respectively. The figure is based on analysis of stable processing intermediates from previous studies ^13,17,43^. (B-D) Northern blots loaded with total RNA from cells induced for Hel66 RNAi for 0, 24 and 48 hours were probed with pre18S (B), ITS2 (C) and ITS3 (D) and for tubulin (loading). The experiment was performed with three different clonal cell lines. One representative northern blot is shown and quantifications of the different precursors (averages of the three clones normalised to tubulin with error bars representing standard error of the mean) are shown below. The 0.61 kb band and its precursor (white frame) are shown with increased contrast underneath the blot (C). The novel processing intermediate of the 5.8 S rRNA is marked with an asterisk. Differences to the uninduced cells were considered statistically significant, when the P-value was smaller than 0.05 (one-way ANOVA and Dunnette’s Post-hoc test; * represents P < 0.05, ** P < 0.01 and *** P < 0.001).

The pre-18S probe detects the initial large 9.2 kb precursor transcript as well as the 3.4 kb and 2.6 kb precursor of the 18S rRNA. We could only detect the 3.4 kb precursor, as the 9.2 kb and the 2.6 kb transcript are unstable ^13,27,43^ and thus not abundant enough for detection (Figure 4B). Upon RNAi depletion of Hel66 there was a 1.5-fold increase in the 3.4 kb precursor within 24 hours of RNAi induction (Figure 4B).

The ITS2 probe detects the initial large 9.2 kb precursor transcript as well as the 5.8 kb cleavage product of the large subunit rRNA precursors and the 0.61 kb precursor of the 5.8S rRNA. Both the 5.8 kb and 0.61 kb precursors were readily detectable on a northern blot, while the 9.2 kb transcript was not abundant enough for detection (see above) (Figure 4C). Upon RNAi depletion of Hel66, we observed a 1.4-fold increase in the 5.8 kb precursor within 24 hours of induction. Interestingly, the amount of 0.61 kb precursor decreased, and instead a slightly larger RNA became visible after 48 hours, indicating that the depletion of Hel66 caused accumulation of a previously unknown precursor of the 0.61 kb intermediate (Figure 4C).

The ITS3 probe detects many intermediates of the six trypanosome rRNAs that are equivalent to the 25/28S rRNA in other eukaryotes. The 5.0 and 5.8 kb intermediates were abundant enough for detection by fluorescent probes (Figure 4D). RNAi depletion of Hel66 caused an increase in the amount of 5.8 kb precursor and a decrease in the amount of 5.0 precursor, indicating an inhibition of the 5.8 to 5.0 processing step.

Taken together, the data indicate an involvement of Hel66 in the processing of ribosomal rRNAs of both ribosomal subunits. At least three different processing steps are inhibited: the processing of the 3.4 and 5.8 kb precursors and the processing of a yet undescribed precursor of the 0.61 kb transcript. This points towards Hel66 having a general function in rRNA processing rather than to it being involved in one particular processing step.

### Loss of Hel66 results in decreased global translation

Any interference with rRNA processing is likely to affect the formation of mature rRNAs and thus protein synthesis. We therefore investigated the effect of Hel66 depletion on translation using the SUnSET assay ^44^. The assay is based on the incorporation of puromycin into growing polypeptide chains. Anti-puromycin antibodies detect puromycin-labelled nascent polypeptides thereby providing information on the rate of global translation. Translation was monitored following RNAi depletion of Hel66. A decrease in global translation to 60% compared to that in uninduced cells (-Tet) was observed 24 h after RNAi induction, with a further decrease to 40% observed after 48 hours of RNAi induction (Figure 5). No signal was detectable without puromycin treatment and a massive decrease in translation to <5% was observed with the translational inhibitor cycloheximide (CHX) (Figure 5). The data show that loss of Hel66 causes a reduction in translation.

**Figure 5:**
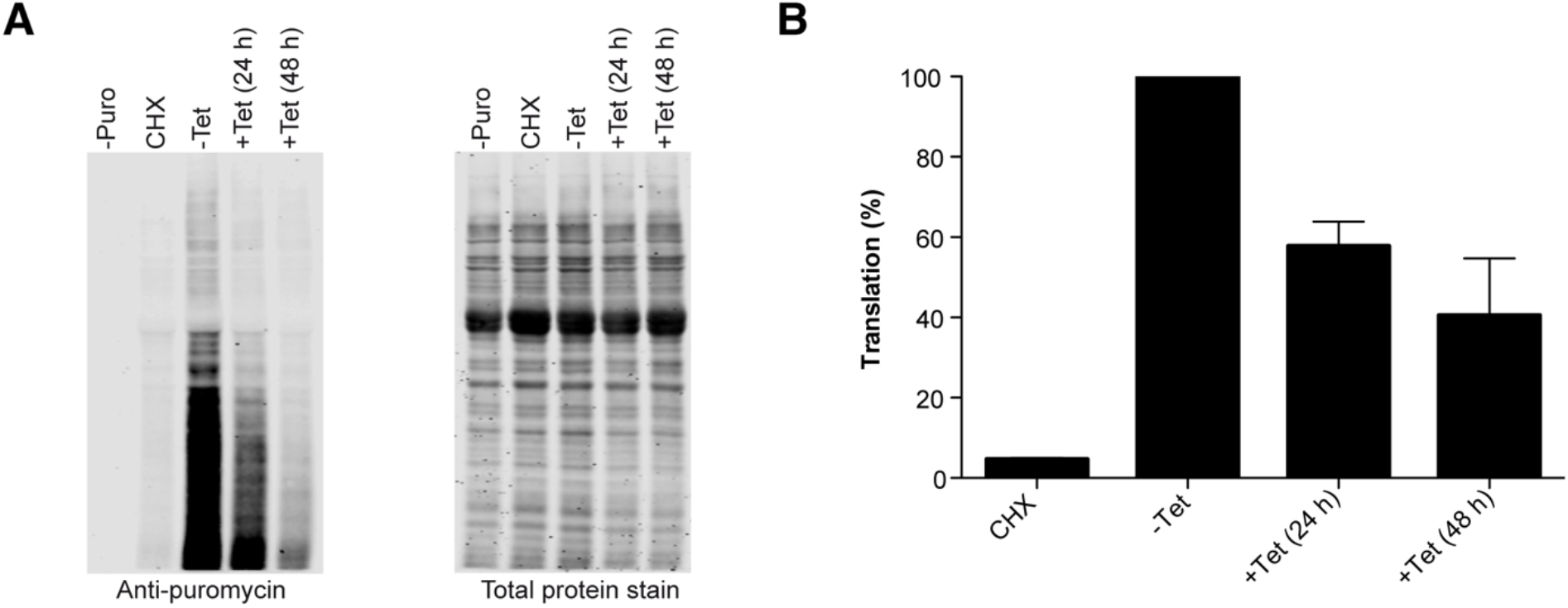
Depletion of Hel66 inhibits protein synthesis. (A) Left: Western blot showing puromycin incorporation into proteins after the SUnSET assay. −Puro = uninduced cells with no puromycin treatment, CHX= uninduced cells treated with cycloheximide (inhibitor of translation) 30 min prior to treatment with puromycin, −Tet = uninduced cells treated with puromycin, +Tet (24 h) = cells treated with puromycin after 24 h of RNAi induction, +Tet (48 h) = cells treated with puromycin after 48 h of RNAi induction. Right: Total protein stain used as a loading control. (B) Quantification of the amount of puromycilated peptides (translation) from the western blot. The experiment was carried out with three different clonal cell lines and the mean values are shown ± SEM.

## DISCUSSION

With Hel66, we have identified and experimentally characterised a DExD/H helicase that is involved in ribosome biogenesis of in *Trypanosoma brucei*. Depletion by RNAi caused a growth and translational arrest as well as an accumulation of precursors of both the large and the small ribosomal subunits, indicative of an essential function of Hel66 in ribosome biogenesis of both ribosomal subunits.

Trypanosomes have about 51 DExD/H helicases ^33,34,37^, and many localise either in nucleolar (19) or nucleoplasmic (9) regions ^39^, indicating that the majority of this protein family may be involved in ribosome processing. A total of 17 of these DExD/H RNA helicases mostly those with nucleolar localisation, have been assigned to yeast homologues with known functions in ribosome biogenesis ^13–15^. These 17 proteins were also co-purified with at least one of four proteins involved in ribosome processing in *Leishmania tarentolae* ^15^. The proteins co-purified with either the SSU protein UTP18 ^45^ or the LSU processome proteomes SSF1, RPL24 and ARX1, that in yeast are involved in early, middle and late processing steps, respectively ^46,47^.

Hel66 is not among the 17 helicases assigned to a yeast homologue ^13–15^. The closest BLAST hits from yeast do not return Hel66 in the *vice versa* BLAST. Hel66 may thus be a trypanosome-unique enzyme, possibly involved in ribosomal processing steps that are absent from yeast and unique to trypanosomes. Hel66 was, however, copurified with SSF1 (Shulamit Michaeli, personal information), confirming its function in early processing of the large ribosomal subunit.

The only other DExD/H RNA helicase that has been identified in the context of rRNA maturation in trypanosomes is Mtr4 ^30^. Mtr4 localises to the nucleoplasm ^39^ and is likely to be part of the TRAMP complex and thus involved in general RNA surveillance in the nucleus together with the nuclear exosome ^30,48^. Depletion of either Mtr4 or exosomal subunits cause accumulation of ribosomal precursor RNAs ^30,31^. In yeast, recent cryoelectron data showed that Mtr4 mediates the direct interaction between the exosome and the 90S pre-ribosomal subunit, indicating a direct function of Mtr4 in rRNA maturation ^49^. At least two trypanosome exosome subunits, Rrp44 and Rrp6 localise to the rim of the nucleolus ^50^, suggesting a similar function of Mtr4 in trypanosomes; Rrp44 may also play an exosome-independent role in early stages of large ribosomal subunit maturation ^32^.

Hel66 depletion resulted mainly in accumulation of the products of the very first pre rRNA cleavage: the 3.4 kb precursor of the 18S rRNA from the SSU and the 5.8 kb precursor of most LSU rRNAs. In addition, there was a decrease in the amount of the 0.61 kb precursor of the 5.8S rRNA while at the same time an rRNA of slightly larger size became visible with the same probe. The likeliest explanation is that the interruption of the processing pathway caused stabilisation of a yet undescribed processing intermediate of the 5.8S rRNA. The data suggest an involvement of Hel66 in rRNA processing pathways of both ribosomal subunits. Given that the precursors differ in stability and only abundant rRNA intermediates were detectable, we cannot define the precise timing of action of Hel66.

The process of rRNA biogenesis is significantly different in trypanosomes which are a rare example for a cell biological model outside the opisthokonts. With Hel66 we have identified an RNA helicase that appears to be unique to trypanosomes and is involved in the processing of rRNA from both ribosomal subunits. This study therefore sheds light on some unique features of the rRNA processing pathways in this early divergent protozoan parasite.

## MATERIALS AND METHODS

### RNA pull-down and mass spectrometry

Hel66 (Tb927.10.1720) was identified from a pulldown assay using the 3’ UTR of VSG121 of *T. brucei.* Briefly, biotinylated RNA containing the first 188 nucleotides of the VSG 3’UTR (including both the 16mer and 8mer motifs) was coupled to streptavidin beads and exposed to *T. brucei* cell lysate. Bound proteins were identified by mass spectrometry. Three different control RNAs were used. Control 1 was the first 188 nucleotides of the VSG 3’UTR with scrambled 16mer and 8mer, control 2 was the reverse complement of control 1 and control 3 was the first 188 nucleotides of the VSG 3’UTR with the 16mer reversed.

### Cloning, expression and purification of recombinant Hel66

The DNA encoding the Hel66 open reading frame (ORF) of the *T. brucei brucei* strain Lister 427 was cloned into the pGEX-4T2 expression vector (GE Healthcare Life Sciences) and the plasmid transfected into ArcticExpress (DE3) *E. coli* cells (Agilent Technologies) for expression of Hel66 fused to an N-terminal GST-tag.

For production of recombinant Hel66 protein, 500 ml Luria-Bertani (LB) medium was inoculated with pre-cultured ArcticExpress (DE3) Hel66 cells. The cells were grown to an OD600 of ~ 0.6, cooled on ice for 30 min, induced with 0.1 mM IPTG and grown overnight at 10 °C. The bacteria were harvested by centrifugation (7,700 × *g*, 5 min, 4 °C), washed once with 100 ml PBS (7,700 × *g*, 5 min, 4 °C) following which the pellet was re-suspended in 50 ml of buffer A (PBS with 10 mM DTT and 1 x protease Inhibitor Cocktail (Roche)). The cells were lysed by sonication with ultrasound (10 sec pulses separated by 1 min breaks) until the solution became transparent. 1% Triton X-100 was added and the lysate was incubated for 30 min at 4°C while rotating. The lysate was centrifuged (15,000 × *g*, 10 min, 4 °C) and the supernatant (clear lysate) was then loaded onto GST GraviTrap columns (GE Healthcare Life Sciences) that had been pre-washed with PBS and buffer B (PBS pH 7.4, 10 mM ATP, 20 mM MgCl_2_, 50 mM KCl, 10 mM DTT, 1 × Protease Inhibitor Cocktail (Roche)). The column was incubated for 30 min with 10 ml of buffer C (PBS pH 7.4, 10 mM DTT, 10 mM ATP, 20 mM MgCl_2_, 40 mM KCl, protease Inhibitor Cocktail) mixed with 23 μl of urea-denatured bacterial lysate, and then washed twice with 10 ml buffer B and twice with 10 ml buffer D (PBS pH 7.4, 5 mM ATP, 20 mM MgCl_2_, 50 mM KCl, 10 mM DTT, 1 x Protease Inhibitor Cocktail (Roche)). The GST-Hel66 protein was eluted with 10 ml elution buffer (50 mM Tris-Cl pH 8.0, 20 mM glutathione) and 2 ml fractions were collected. Fractions containing GST-Hel66 were identified by SDS-PAGE, pooled, concentrated and the buffer exchanged to PBS using an Amicon ultra-15 10K centrifugal filter (Merk-Millipore). The protein was frozen in liquid nitrogen and stored at −80 °C.

### Trypanosome cell lines and culture conditions

*T. brucei* 13-90 cells ^51^ and 2T1 cells (both modified monomorphic bloodstream form *Trypanosoma brucei brucei* strain 427, variant MITat1.2) ^52^ were used throughout this study. All transgenic cell lines generated in this study are based on the 2T1 cell line. Cells were grown in HMI-9 medium ^53^ supplemented with 10% fetal calf serum (FCS) (Sigma-Aldrich, St. Louis, USA) and incubated at 37°C and 5% CO_2_. For maintenance of previously transfected plasmids, *T. brucei* 13-90 cells were cultured with 5 μg/ml hygromycin and 2.5 μg/ml G418 whereas *T. brucei* 2T1 cells were cultured with 0.1 μg/ml puromycin and 2.5 μg/ml phleomycin. Cell numbers were monitored using either a haemocytometer or a Z2 Coulter counter (Beckman Coulter).

### Plasmid construction and generation of transgenic trypanosome cell lines

The Hel66 RNAi cell line was generated using the pGL2084 vector ^54^ and *T. brucei* 2T1 cells. The conserved nature of the DExD/H protein family prevented the usage of the full open reading frame sequence of TbDHel116 for RNAi. Instead, two short fragments from the non-conserved N- and C-terminal extension regions of Hel66 were amplified with primer pairs MBS37/MBS38 and MBS39/MBS40, respectively, and then joined together in an additional PCR step using the primer pair MBS37/MBS40 to obtain a 305 bp DNA fragment with AttB adaptor sequences. This DNA fragment was cloned into pGL2084 by a BP Recombinase reaction (Invitrogen) following the manufacturer’s instructions to generate pGL2084_Hel66. The plasmid was linearised with NotI, transfected into *T. brucei* 2T1 cells and positive transfectants were selected with 2.5 μg/ml hygromycin. RNAi was induced with 1 μg/ml tetracycline.

For C-terminal tagging of one endogenous Hel66 allele with HA or mNeonGreen, the PCR based tagging system using pMOTag3H or pMOTag_mNG (a modified pMOTag3G plasmid with the GFP replaced by mNeonGreen) as templates, respectively, was used ^55^. All primers used for plasmid construction and generation of transgenic cell lines are provided in Table S1.

All transfections for generation of transgenic cell lines were carried out with 3 × 10^7^ trypanosome cells, electroporated with 10 μg of linearised plasmid DNA or PCR product using the Amaxa Basic Parasite Nucleofector Kit 1 and Nucleofector II device (Lonza, Switzerland, program X-001).

### Western blot

Whole cell protein lysate from 1 × 10^6^ trypanosome cells was separated on 12.5% sodium dodecyl sulphate (SDS)-polyacrylamide gels and transferred onto nitrocellulose membranes (GE Healthcare Life Sciences). The membranes were blocked by incubation with 5% milk powder in PBS for 1 h at room temperature (RT) or overnight at 4 °C. Primary antibodies (rabbit anti-VSG221, 1:5000) and mouse anti-PFR antibody (L13D6, 1:20)) ^56^ were then applied in PBS/1% milk/0.1% Tween-20 solution for 1 h at RT. After four washes (5 min each) with PBS/0.2% Tween-20, IRDye 800CW-conjugated goat-anti-rabbit and IRDye 680LT-conjugated goat-anti-mouse secondary antibodies were applied in PBS/1% milk/0.1% Tween-20 solution for 1 h at RT in the dark. The membranes were washed four times (5 min each in the dark) with PBS/0.2% Tween-20 followed by a final 5 min wash with PBS. Blots were analysed using a LI-COR Odyssey system (LI-COR Biosciences).

### RNA extraction and Northern blot analysis

Total RNA was extracted from 1 × 10^8^ trypanosome cells using the Qiagen RNeasy Mini Kit (Qiagen, Netherlands) following the manufacturer’s instructions. Northern blot analyses were carried out using 8 μg of total RNA. The RNA was denatured with glyoxal at 50°C for 40 min as previously described ^57^ and loaded on a 1.5% agarose gel containing 10 mM sodium phosphate, pH 6.9. The RNA was transferred overnight to a Hybond N+ nylon membrane (GE Healthcare) by upward capillary transfer. After transfer, the RNA was UV crosslinked (1200×100 μJ/cm^2^) onto the membranes and deglyoxylated by baking at 80 °C for 1 h. The membranes were then prehybridised at 42°C for 1 h in hybridisation solution (5x SSC (3 M NaCl, 0.3 M tri-sodium citrate, pH 7.0), 10% 50x Denhardt’s solution (1% BSA, 1% polyvinylpyrrolidone, 1% Ficoll), 0.1% SDS, 100 μg/ml heparin, 4 mM tetrasodium pyrophosphate). Membranes were hybridised overnight at 42 °C with hybridisation solution containing the fluorescently labelled oligonucleotide probes (10 nm each) (listed in Table S1). After hybridisation, the membranes were washed three times for 10 min with northern wash buffer (2x SSC buffer, 0.1% SDS), dried and the blots analysed using a LI-COR Odyssey system (LI-COR Biosciences).

### Quantification of *VSG* mRNA

*VSG* mRNA was quantified using RNA dot blots as previously described ^58^. Briefly, 3 μg of glyoxal-denatured RNA was applied to a nitrocellulose membrane (Hybond-N) using a Minifold Dotblotter (Schleicher & Schuell, Germany). The blots were hybridised over night at 42 °C with a *VSG221* oligonucleotide probe coupled to IRDye 682 and a *tubulin* oligonucleotide probe coupled to IRDye 782, which was used as a loading control. Blots were analysed using the LI-COR Odyssey system (LI-COR Biosciences).

### Translation assay

The SUnSET (Surface Sensing of Translation) assay was used to monitor global translation ^59^. 1 × 10^7^ mid-log phase trypanosome cells were treated with 10 μg/ml puromycin for 30 min, washed once with trypanosome dilution buffer (TDB; 5 mM KCl, 80 mM NaCl, 1 mM MgSO_4_, 20 mM Na_2_HPO_4_, 2 mM NaH_2_PO_4_, 20 mM glucose, pH 7.6) and boiled in 1x protein sample buffer (2% SDS, 10% glycerol, 60 mM Tris–HCl, pH 6.8, 1% β-mercaptoethanol) (5 min, 100 °C). As controls, cells were either treated with 50 μg/ml of the translational inhibitor cycloheximide for 30 min prior to treatment with puromycin, or not treated with puromycin (negative control). The protein samples (containing 1 × 10^6^ cell equivalents) were resolved on a 12.5% SDS gel and puromycin-labelled peptides were detected with anti-puromycin (1:5000 mouse anti-puromycin, clone 12D10; Sigma). Prior to antibody detection, REVERT 700 total protein stain (LI-COR Biosciences) was carried out according to the manufacturer’s instructions and used as a loading control.

### RNA electrophoretic mobility shift assay (REMSA)

REMSA experiments were carried out with the Light Shift Chemiluminescent RNA EMSA kit (Thermo Fisher Scientific) according to the manufacturer’s instructions. GST-Hel66 was incubated in reaction buffer (1x REMSA buffer, 5% glycerol, 0.1 μg/μl tRNA) containing 10 nM biotin-labelled 16mer RNA probe (Table S1) for 30 min at room temperature. For competition assays, unlabelled 16mer RNA or Biotin-IRE control RNA (RNA encoding the iron response element, provided by the kit) were added in 200-fold excess. Samples were applied to a 5% native polyacrylamide gel in 0.5x TBE (Tris-borate-EDTA) buffer. After transfer to a Hybond N+ nylon membrane (GE Healthcare), samples were UV cross-linked and the biotin signal detected with HRP–conjugated streptavidin using the Chemiluminescent Nucleic Acid Detection Module (Thermo Fisher Scientific) according to the manufacturer’s instructions.

### Microscopy

1 × 10^7^ cells were harvested by centrifugation at 1400 × *g* for 10 min. The cells were washed with PBS and fixed overnight in 4% formaldehyde and 0.05% glutaraldehyde at 4 °C. Fixed cells were washed twice with PBS and mounted in Vectashield mounting medium with DAPI (Vector Laboratories Inc.). Images were acquired with an automated DMI6000B wide field fluorescence microscope (Leica microsystems, Germany), equipped with a DFC365FX camera (pixel size 6.45 μm) and a 100x oil objective (NA 1.4). Fluorescent images were deconvolved using Huygens Essential software (Scientific Volume Imaging B. V., Hilversum, The Netherlands) and are presented as Z-projections (method sum of slices). In order to visualise the localisation of Hel66, the DAPI signal is shown in magenta and the protein is shown in green. Image analysis was carried out using the Fiji software ^60^.

## ACKNOWLEDGEMENTS

We thank Henriette Zimmermann, Kathrin Weißenberg, Alyssa Borges and Paula Castaneda Londono for technical assistance. MB-S was supported by a grant from the German Excellence Initiative awarded to the Graduate School of Life Sciences, University of Würzburg. M.E. is funded by the DFG GRK 2157, DFG grants EN305, SPP1726, GIF grant I-473-416.13/2018, the EU ITN Physics of Motility, and the BMBF NUM Organo-Strat. M.E. is a member of the Wilhelm Conrad Röntgen Centre for Complex Material Systems (RCCM).

## AUTHOR CONTRIBUTIONS

M.B-S., N.G.J., S.K., F.B., and M.E. conceived and designed the experiments; M.B-S., N.J.I. and M.S. conducted the experiments; M.B-S., N.G.J., M.S., F.B., S.K. and M.E. analysed and interpreted the results; M.B-S., N.G.J., S.K. and M.E. wrote the initial manuscript. All the authors reviewed the manuscript.

## SUPPLEMENTARY INFORMATION

**Supplementary Figure S1:**
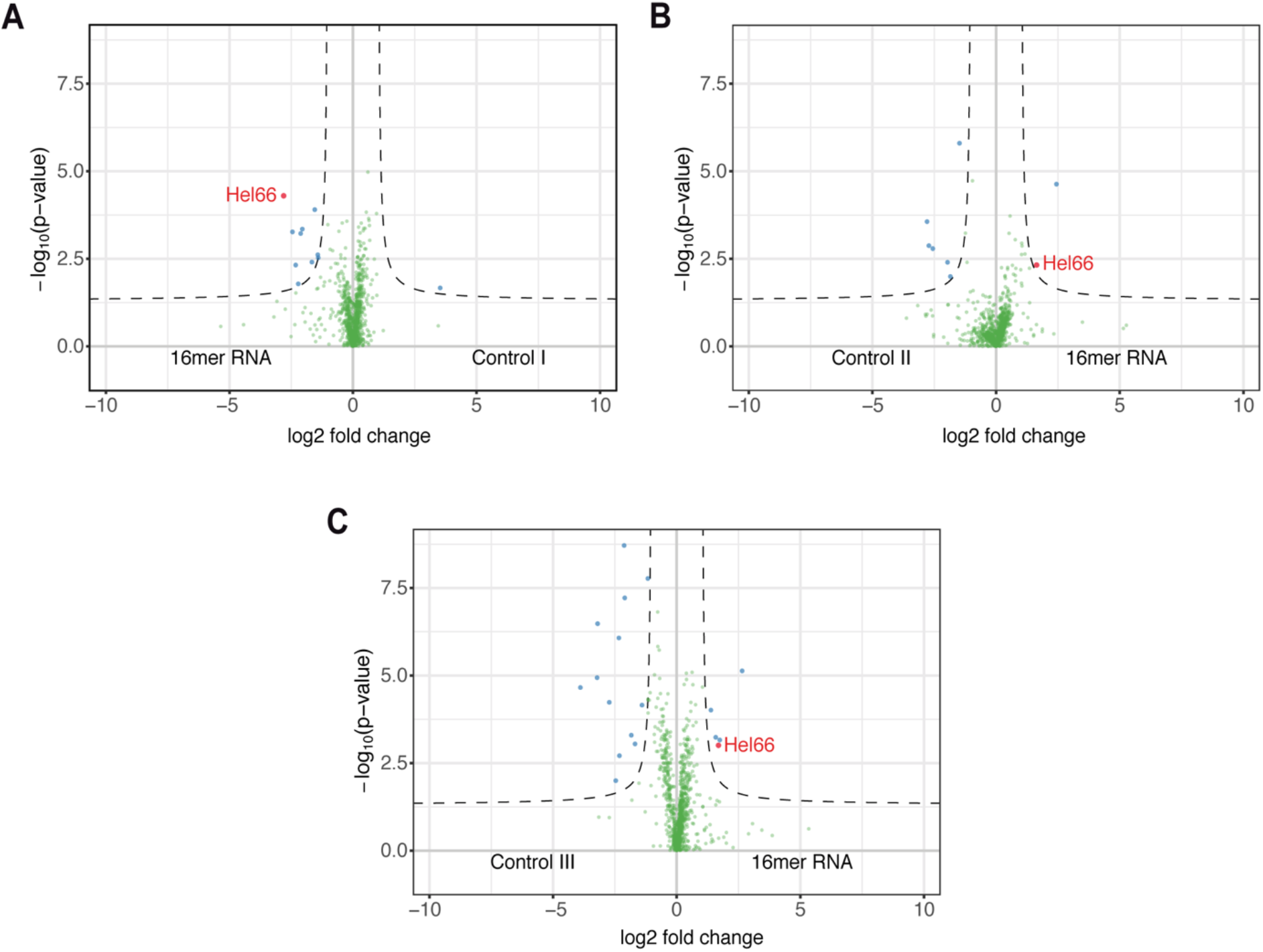
Volcano plots showing potential interaction partners of the 16mer motif. The first 188 nucleotides of the VSG121 3’ UTR (following the stop codon) harbouring the 16mer and 8mer motifs were in vitro transcribed and used as bait. (A-C) Triplicate assays using different controls (control I: first 188 nucleotides of the VSG121 3’ UTR with scrambled 16mer and 8mer, control II: reverse complement of control I, control III: first 188 nucleotides of the VSG121 3’ UTR with reverse complement of 16mer).

**Supplementary Figure S2:**
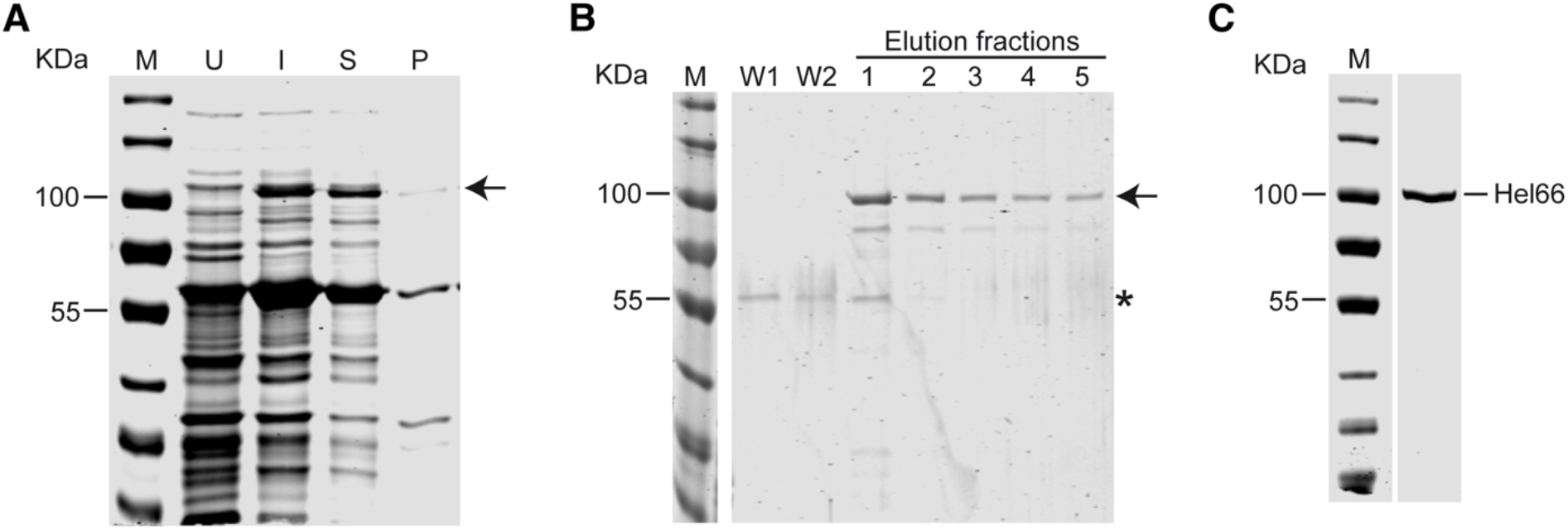
Expression and purification of recombinant GST-tagged Hel66 (GST-Hel66). (A) SDS page gel showing GST-Hel66 (arrow) expressed in ArcticExpress *E. coli* cells. M = Marker, U = uninduced, I = induced, S = supernatant (soluble fraction) and P = pellet (insoluble fraction) (B) Purification of GST-Hel66 (arrow) using GST-Gravitrap column. M = Marker, W1= flow-through from first wash, W2 = flow-through from second wash, 1-5 different elution fractions. Asterisk (*) indicates the position of the co-purifying chaperon Cpn60 of 55 kDa molecular weight. (C) Concentrated purified GST-Hel66 protein.

**Supplementary Figure S3:**
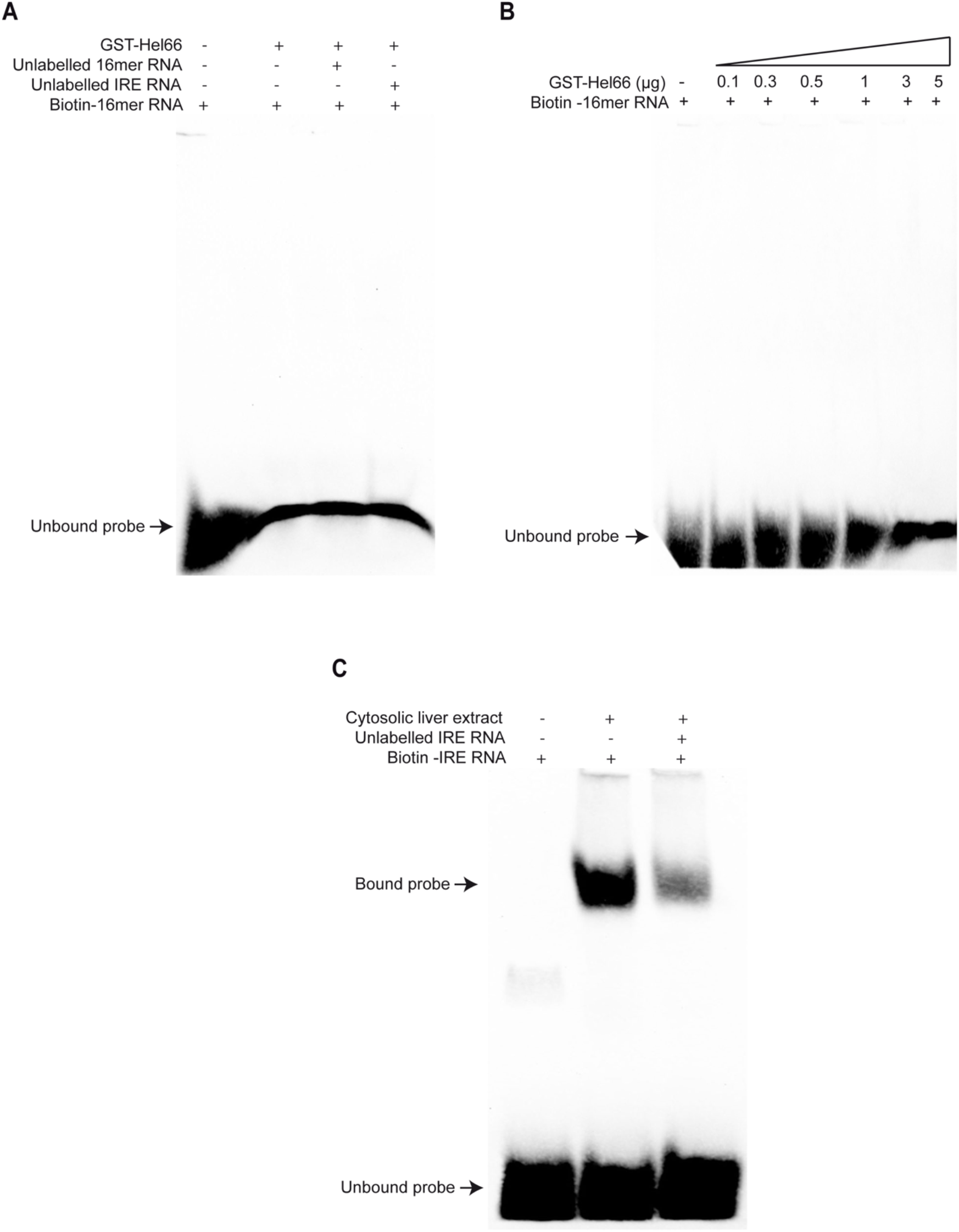
The 16mer RNA does not interact with GST-Hel66. (A) REMSA using 16mer RNA and GST-Hel66. 10 nM of biotinylated 16mer RNA was incubated at room temperature with 5 μg of GST-Hel66 in a reaction mix of 20 μl for 30 min. Competition assays were carried out by adding 200-fold excess of either unlabelled 16mer RNA or an unrelated RNA (IRE RNA). (B) REMSA assay with the 16mer RNA and different concentrations of GST-Hel66 protein. 10 nM of biotinylated 16mer RNA was incubated at room temperature with GST-Hel66 in a reaction mix of 20 μl for 30 min. (C) Positive control from the REMSA kit showing functional binding/interaction between IRE (iron-response element) RNA and IRP (iron-response protein).

**Supplementary Figure S4:**
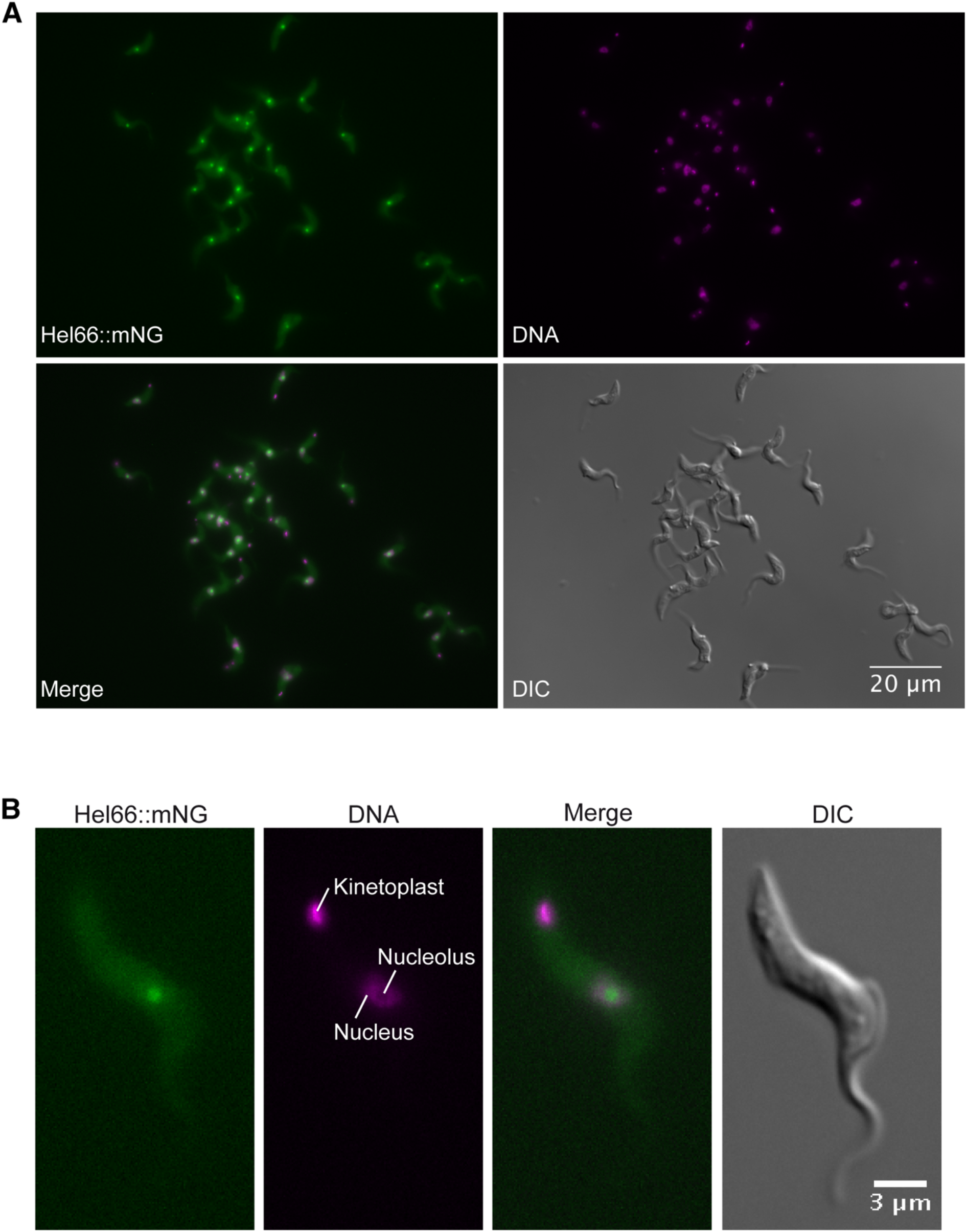
Hel66 fused to mNeonGreen localises to the nucleolus in *T. brucei* bloodstream form cells. (A) Localisation of Hel66 in a population of cells. (B) Raw images of one example of a cell showing the nucleolar localisation of Hel66.

**Supplementary Figure S5:**
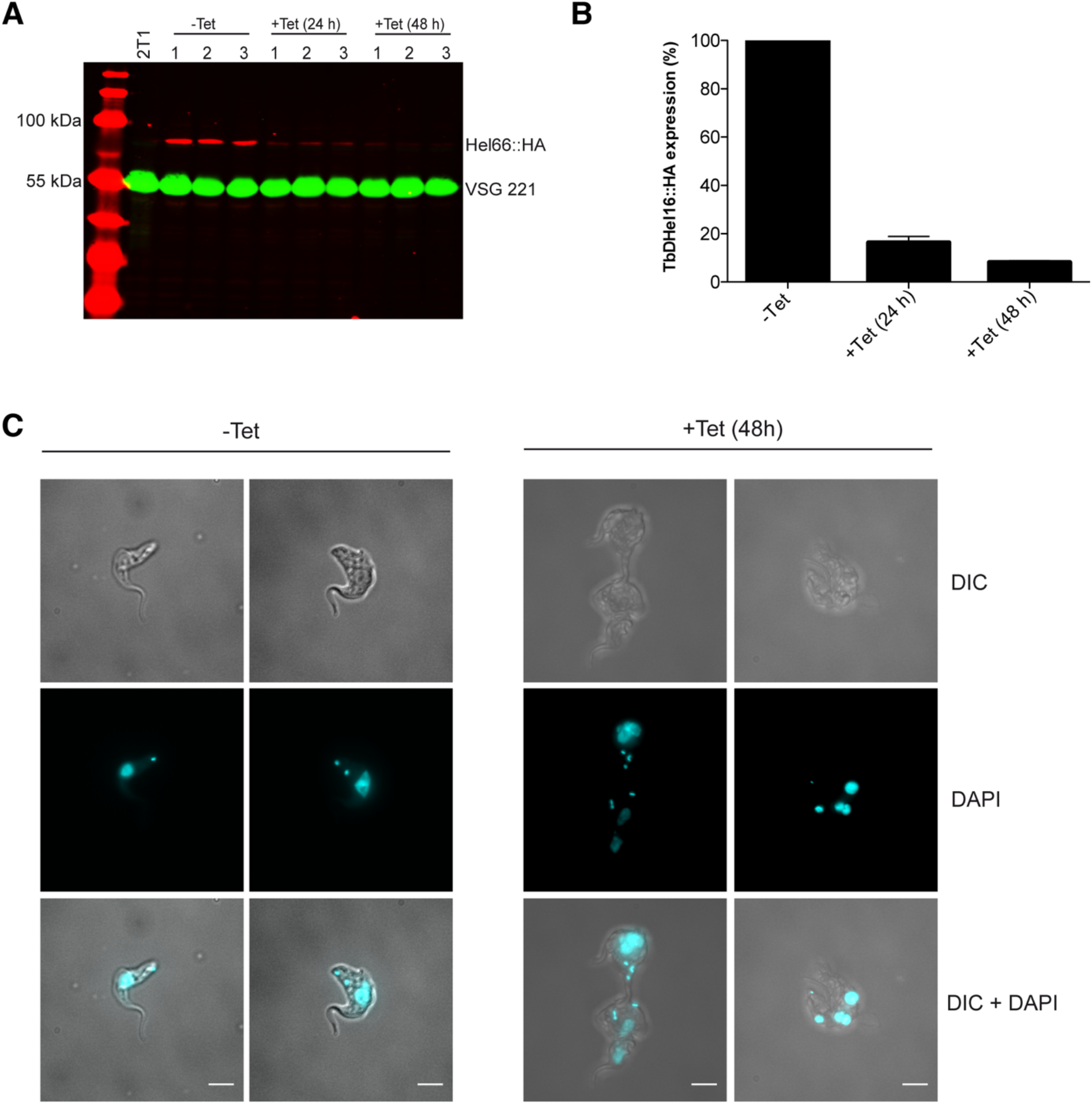
RNAi-mediated depletion of Hel66. (A) Western blot showing the protein amounts of Hel66::HA for three independent clonal cell lines over a time following RNAi induction. 2T1 is the parental cell line. VSG221 served as a loading control (B) Quantification of the protein amounts of Hel66::HA. The signal intensity of Hel66::HA was normalised to the VSG221 signal. Average data from the three independent clonal cell lines are shown, with the standard error of the mean (SEM) presented by error bars. (C) Examples of cells with aberrant morphology following RNAi depletion of Hel66 (right); uninduced cells are shown as a control (left). Scale bar = 5μm.

**Supplementary Figure S6.**
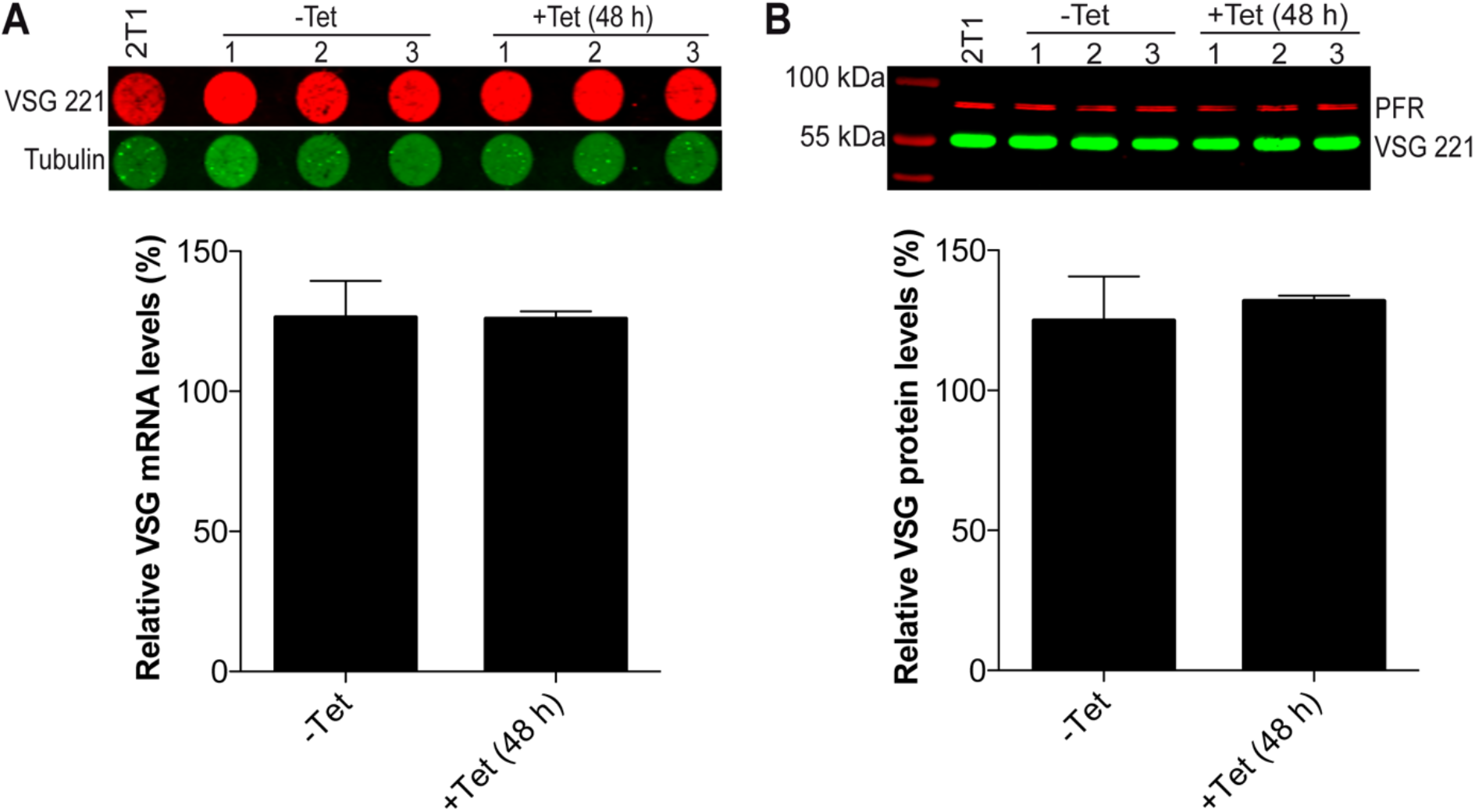
*VSG221* mRNA and protein levels upon depletion of Hel66. (A) *VSG221* mRNA levels in uninduced (-Tet) and induced (+Tet 48 h) Hel66-RNAi cells. *Tubulin* mRNA was used as a loading control. Average data of three independent clonal cell lines are shown with error bars representing the standard error of the mean (SEM). (B) VSG221 protein levels in uninduced (-Tet) and induced (+Tet 48 h) Hel66-RNAi cells. A paraflagellar rod protein (PFR) was used as a loading control. Average data of three independent clonal cell lines are shown with error bars representing the standard error of the mean (SEM).

**Supplementary Table S1.**
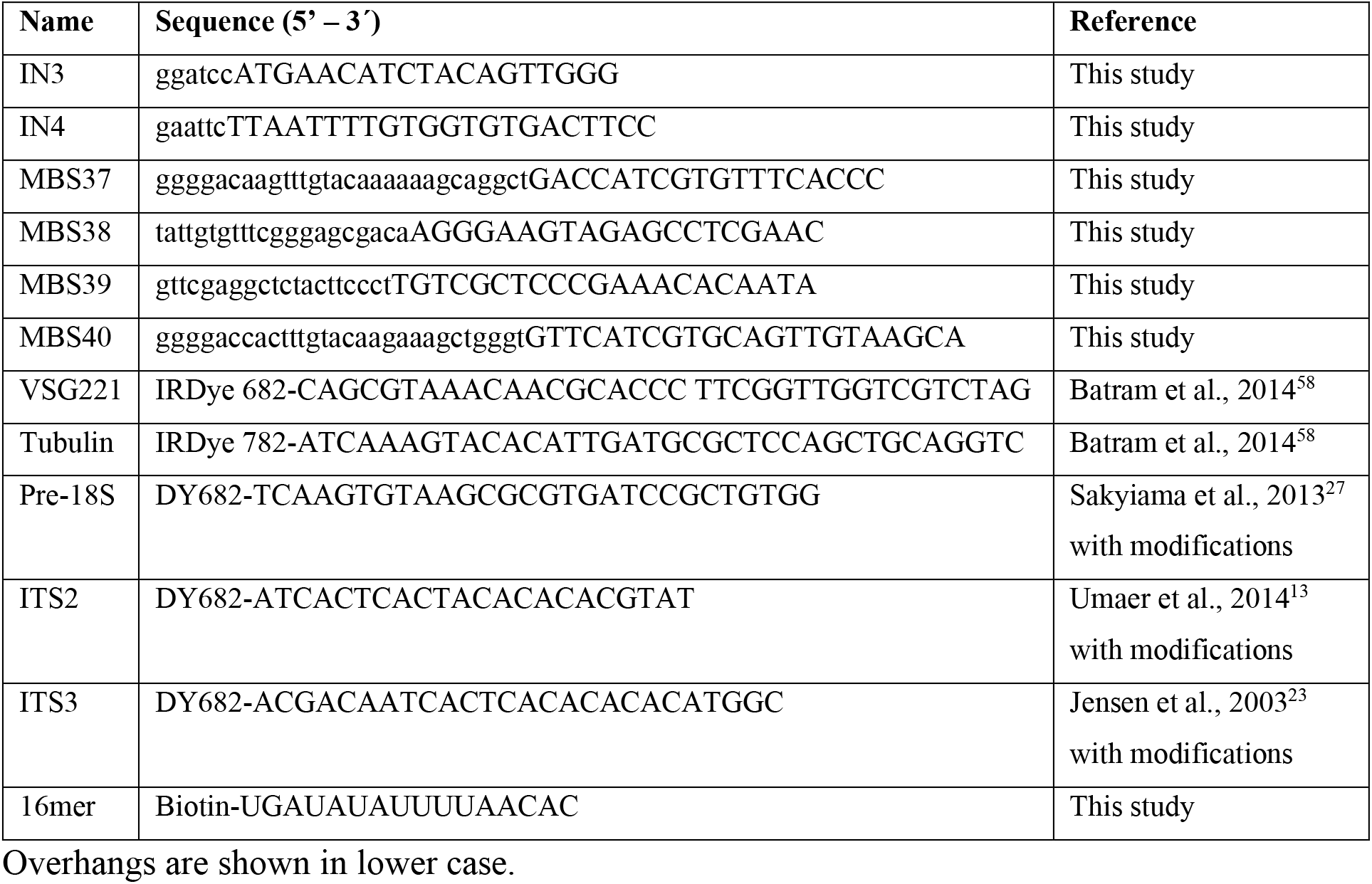
List of primers and probes used in the study

